# Hierarchical Bayesian Modelling of Interoceptive Psychophysics

**DOI:** 10.1101/2025.08.27.672360

**Authors:** Arthur S. Courtin, Jesper Fischer Ehmsen, Leah Banellis, Francesca Fardo, Micah G. Allen

## Abstract

Interoception, the capacity to sense, perceive, and metacognitively appraise viscerosensory and homeostatic signals, is a growing focus in psychology and psychiatry. Adaptive psychophysical tasks now allow quantification of perceptual sensitivity, bias, and precision in cardiac and respiratory domains. However, accurately estimating these parameters often requires large numbers of trials or participants, posing practical challenges, especially in clinical research where participant availability and tolerance are limited. One approach to reduce participant burden while maintaining statistical rigour is to optimise data analysis. Here, we present hierarchical Bayesian models tailored for cardiac and respiratory interoceptive psychophysics that efficiently estimate sensitivity, bias, and precision at both individual and group levels. Using simulations and empirical data, we validate these models and demonstrate that they allow enhanced inference relative to conventional approaches. To support adoption, we provide openly-accessible resources, including a tutorial on how to implement and fit these models in R (written with researchers without modelling expertise in mind) and an app for sample-size justification. These tools facilitate robust, efficient, and generalisable modelling of interoceptive performance, enabling rigorous studies even with limited trials or participant availability.

## Introduction

Interoception, the sensing and appraisal of internal bodily states, has emerged as a key focus in psychology and psychiatry, linking brain-body signalling to cognition, emotion, and self-awareness (Cameron, 2001; Craig, 2002). For example, interoceptive monitoring of cardiac signals has been linked to emotional arousal and fear processing, and it is frequently investigated in relation to anxiety, panic, and psychosis (Adams et al., 2022; Ehlers et al., 1995; Jeganathan et al., 2025). Despite this potential as a psychiatric biomarker, methods for quantifying and modelling interoception remain limited. To address this issue, we present a hierarchical Bayesian modelling approach for interoceptive psychophysics, spanning the respiratory and cardiac domains.

Interoception research faces several methodological limitations (Desmedt et al., 2023). In particular, commonly used methods such as the heartbeat counting task or self-report questionnaires suffer from psychometric issues related to construct validity and other related issues (Desmedt, Heeren, et al., 2022; Desmedt, Van Den Houte, et al., 2022; Zamariola et al., 2018). Classical psychophysical tasks address some of these limitations, but require a substantial number of trials for reliable estimates, resulting in long testing times and high cognitive burdens that are often impractical for paediatric or clinical populations. Even in healthy volunteers, extended testing sessions risk reducing data quality due to distraction, habituation, or fatigue. This trade-off between validity and feasibility indicates the need for methods that yield valid and reliable estimates while minimising burden on participants. Without such advances, rigorous investigation of the proposed link between interoception and mental health will remain substantially constrained.

To this end, we recently developed and validated two new tasks: the Heart Rate Discrimination Task (HRDT)(Legrand et al., 2022) and the Respiratory Resistance Sensitivity Task (RRST)(Nikolova et al., 2022). These tasks enable the estimation of individual participants’ cardiac and respiratory interoceptive psychometric functions, whose thresholds and slopes provide indices of perceptual sensitivity or bias and perceptual precision, respectively. Unlike classical tasks, the HRDT and RRST use an adaptive Bayesian algorithm to select stimulus intensity at each trial based on the participant’s previous responses, drastically reducing the number of trials required to obtain reliable estimates (Kontsevich & Tyler, 1999). Adoption of these tasks partially helps resolve the trade-off, improving both theoretical validity and estimation efficiency.

Beyond adaptive stimulus placement, the number of trials needed to reach a specific level of estimation precision can also be reduced by analysing the data more efficiently. In most psychophysical research (including in the exteroceptive domain), adaptive algorithms are used both to choose the stimuli and to estimate each participant’s threshold and slope. These values are then treated as single data points in the group analysis. This procedure is limited in two ways: it ignores the uncertainty associated with each individual estimate and it prevents partial pooling of information across participants (Gelman, 2006). As a result, group-level parameters can be biased, variability estimates corrupted, and statistical power reduced, especially when only a few trials are used or when data is noisy.

Following others (Mezzetti et al., 2023; Prins, 2024), we use an alternative approach: rather than analysing the parameter estimates returned by the Bayesian adaptive algorithm, we fit a hierarchical Bayesian model directly to the raw trial-wise data (stimulus intensity and participant response). This model simultaneously estimates individual-and group-level psychometric function parameters, explicitly accounting for the nested structure of the data (e.g., trials are contained with participants, themselves within groups) and enabling principled pooling of information across participants. Compared with conventional methods, this should yield more precise group-level estimates, offering equivalent statistical power with fewer trials.

To evaluate the validity and practical benefits of this approach, we conducted a series of analyses on simulated and empirical data. First, we built hierarchical models tailored to our tasks and assessed their ability to recover true parameter values (parameter recovery) and to distinguish between alternative data-generating processes (model recovery). Second, we fit these models to a large existing dataset to derive empirically informed priors and establish a realistic generative basis for simulations. Third, we used simulations to systematically compare hierarchical and conventional analyses across a range of design choices (trial number, sample size, and inference on threshold or slope). This allowed us both to quantify the power gains afforded by hierarchical modelling and to provide empirically grounded guidance on the minimal testing burden required for rigorous inference, regardless of the analytical approach. Finally, we provide a documented toolkit of open resources to facilitate the adoption of hierarchical models by researchers working with the HRDT and RRST, regardless of their prior modelling experience. These resources include a Shiny app for exploring the impact of design choices on power to aid sample size selection and justification, ready-made Stan models for common experimental designs, and an *R* script demonstrating how to use the *brms* library to fit these hierarchical models to data, diagnose them, as well as interpret and report their results (Bürkner, 2017; R Core Team, 2021).

## Methods

The simulations and analyses reported in this manuscript were conducted in MATLAB (The Mathworks, USA) and R (R Core Team, 2021), using the Rstudio interface (Allaire, 2012), and toolboxes *Palamedes* (Prins & Kingdom, 2018), *cmdstanr* (Gabry et al., 2024), *loo* (Vehtari et al., 2024), and *tidyverse* (Wickham et al., 2019).

### Tasks

#### The Heart Rate Discrimination Task

The Heart Rate Discrimination Task (HRDT) is a psychophysical paradigm designed to measure the bias and precision of individuals’ beliefs about their own cardiac signals (c.f. Legrand *et al*. for an in-depth description). Participants begin each trial by attending to their heart for a brief period, during which their actual heart rate is measured via pulse oximetry. This is followed by the presentation of auditory tones paced at a frequency slightly above or below their recorded heart rate. Participants are asked to judge whether the tone sequence was faster or slower than their perceived heart rate during the preceding interval, making a two-alternative forced choice. After making this binary decision, they provide a confidence rating on a visual analogue scale ranging from ‘guess’ to ‘certain’.

To account for potential confounds such as general temporal estimation ability, the task includes an exteroceptive control condition in which participants make the same faster/slower judgement, but between two sequences of tones unrelated to their own heart rate.

#### The Respiratory Resistance Sensitivity Task

The Respiratory Resistance Sensitivity Task (RRST) is a psychophysical paradigm designed to measure sensitivity and precision of inspiratory resistance detection (c.f. Nikolova *et al*. for an in-depth description). On each trial, participants take two successive inhalations through a closed respiratory circuit. During one of these inhalations, a brief increase in resistance is introduced by partially compressing the circuit tubing using a computer-controlled motor. Following the pair of breaths, participants indicate which inhalation felt more difficult, making a two-interval forced-choice judgement. They then rate their confidence in this decision using a visual analogue scale ranging from ‘guess’ to ‘certain’.

#### Adaptive stimulus selection

Rather than using a pre selected or random sequence of stimulus intensities, the HRDT and RRST employ an adaptive Bayesian thresholding algorithm known as the □ (psi) method to optimise stimulus selection based on participants’ previous responses (Kontsevich & Tyler, 1999). The method is initialised with vectors containing possible values for the threshold and the slope of the psychometric function (PF), as well as a vector of potential stimulus intensities. From these vectors, a grid is constructed that represents the approximate joint probability distribution of the threshold and slope parameters.

At the start of each trial, this distribution is used to estimate the expected entropy reduction associated with presenting each possible stimulus intensity, which corresponds to the expected information gain about the PF parameters. The stimulus intensity predicted to yield the greatest reduction in entropy is then selected and presented to the participant. Following the participant’s response, the joint probability distribution of the threshold and slope parameters is updated accordingly.

### Hierarchical modelling

The □ method also returns estimates for the slope and threshold of the PF, derived from the probability distribution it tracks across trials. However, using these point estimates as data for group-level inference is problematic because it ignores the uncertainty of each individual estimate and prevents information from being shared across participants. To address this, we propose to use hierarchical PF models to fit the trial-wise data generated by the tasks, enabling joint estimation of parameters across individuals and conditions. This approach preserves uncertainty, enables principled pooling of information, and improves the robustness of group-level inference.

#### HRDT model parameterisation

In the HRDT, the PF describes how the probability of judging a tone sequence as “faster” depends on the difference in beats per minute (ΔBPM) between the auditory feedback and either the participant’s own heart rate (interoceptive condition) or a reference tone (exteroceptive condition). This relationship is modelled by a function with three parameters: the threshold (α), the slope (β), and the lapse rate (λ) (Fig. 1).

**Figure 1.**
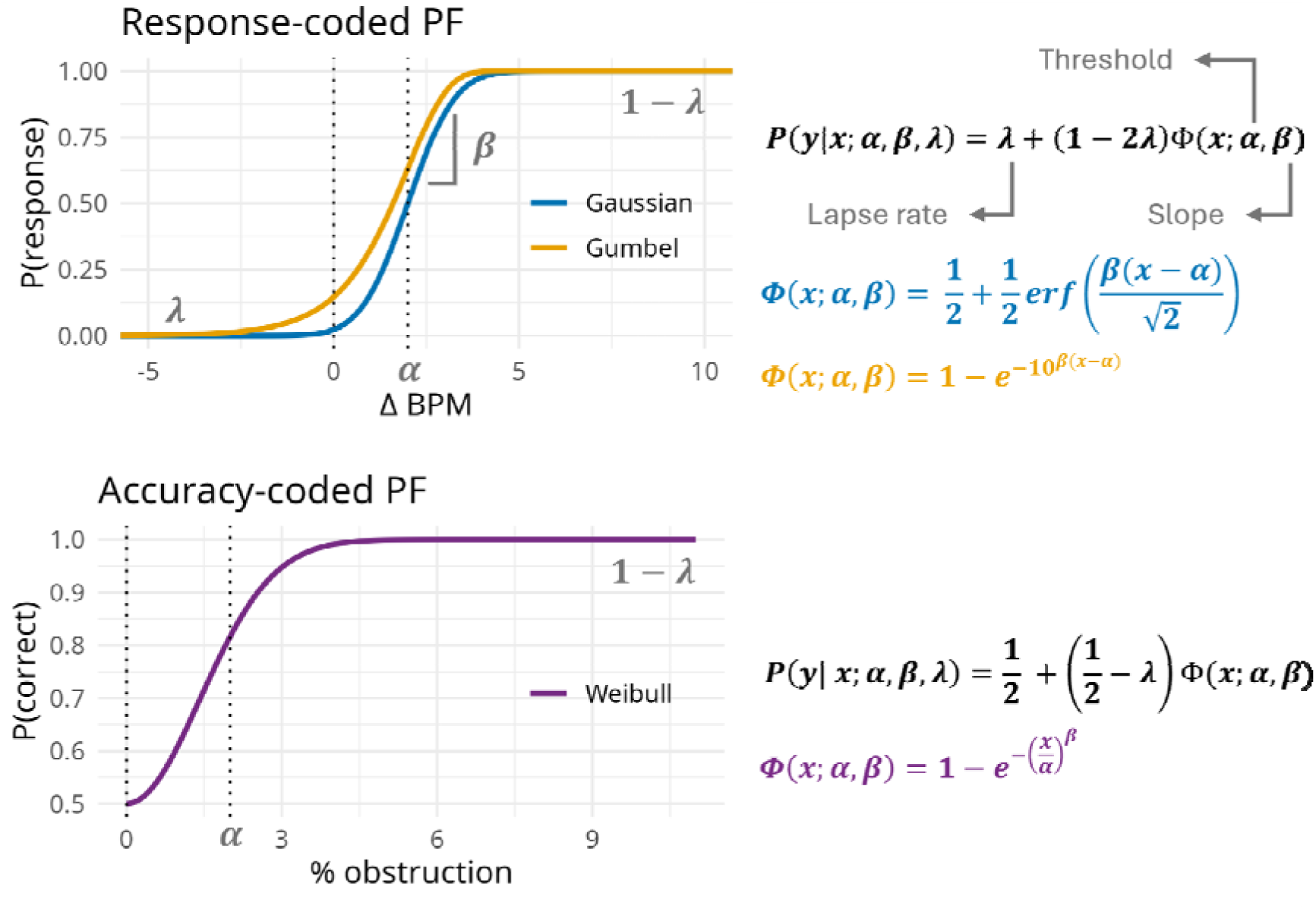
Psychometric function formulations. Top panel: Psychometric functions (PF) used for the Heart Rate Discrimination Task (HRDT). The plot shows two possible PF formulations (i.e, the Gaussian and the Gumbel) relating ΔBPM to the probability of a “faster” response. The threshold (α) sets the horizontal position of the function, the slope (β) controls its steepness, and the lapse rate (λ) adjusts the upper and lower asymptotes. The Gaussian is symmetrical around the threshold, whereas the Gumbel is left skewed. Bottom panel: PF used for the Respiratory Resistance Sensitivity Task (RRST). This Weibull function models the probability of a correct response as a function of relative obstruction. The minimum probability taken by the function is fixed at chance level (50%) and the lapse rate defines the maximum probability. Parameter α corresponds to the occlusion level leading to ∼80% accuracy (assuming L=0). Parameters α and β jointly determine the shape of the PF. The equations for all PFs are shown on the right.

Different mathematical formulations can be used for the PF (Kingdom & Prins, 2010). Prior work with the HRDT used a Gaussian cumulative distribution function (CDF), which is symmetrical around the threshold. However, if responses to positive versus negative ΔBPM differ in shape, a left-skewed function, such as the Gumbel CDF, may provide a better fit. We therefore compare the Gaussian and the Gumbel formulations in our analyses.

In the case of the Gaussian PF, the threshold α represents the ΔBPM value at which participants are equally likely to respond “faster” or “slower” (i.e., the point of subjective equality or bias). In the case of the Gumbel, α corresponds to the ΔBPM value where the probability of response being “faster” is ∼63% (1-exp(-1)), assuming a lapse rate of zero. The slope β controls how quickly the probability of a “faster” response changes as the ΔBPM increases, providing an index of precision. The lapse rate λ accounts for incorrect responses due to inattentiveness or random responses and determines how much the lower and upper bounds of the function deviate from zero and one.

For each PF formulation, we implemented two hierarchical models in Stan (Stan Development Team, 2019): one for single-condition data (interoceptive or exteroceptive only) and one for data including two conditions (e.g., the interoceptive and exteroceptive HRDT conditions). The models estimate participant-level threshold, slope, and lapse rate parameters, along with the mean and standard deviations of their group-level distributions (Fig. 2).

**Figure 2.**
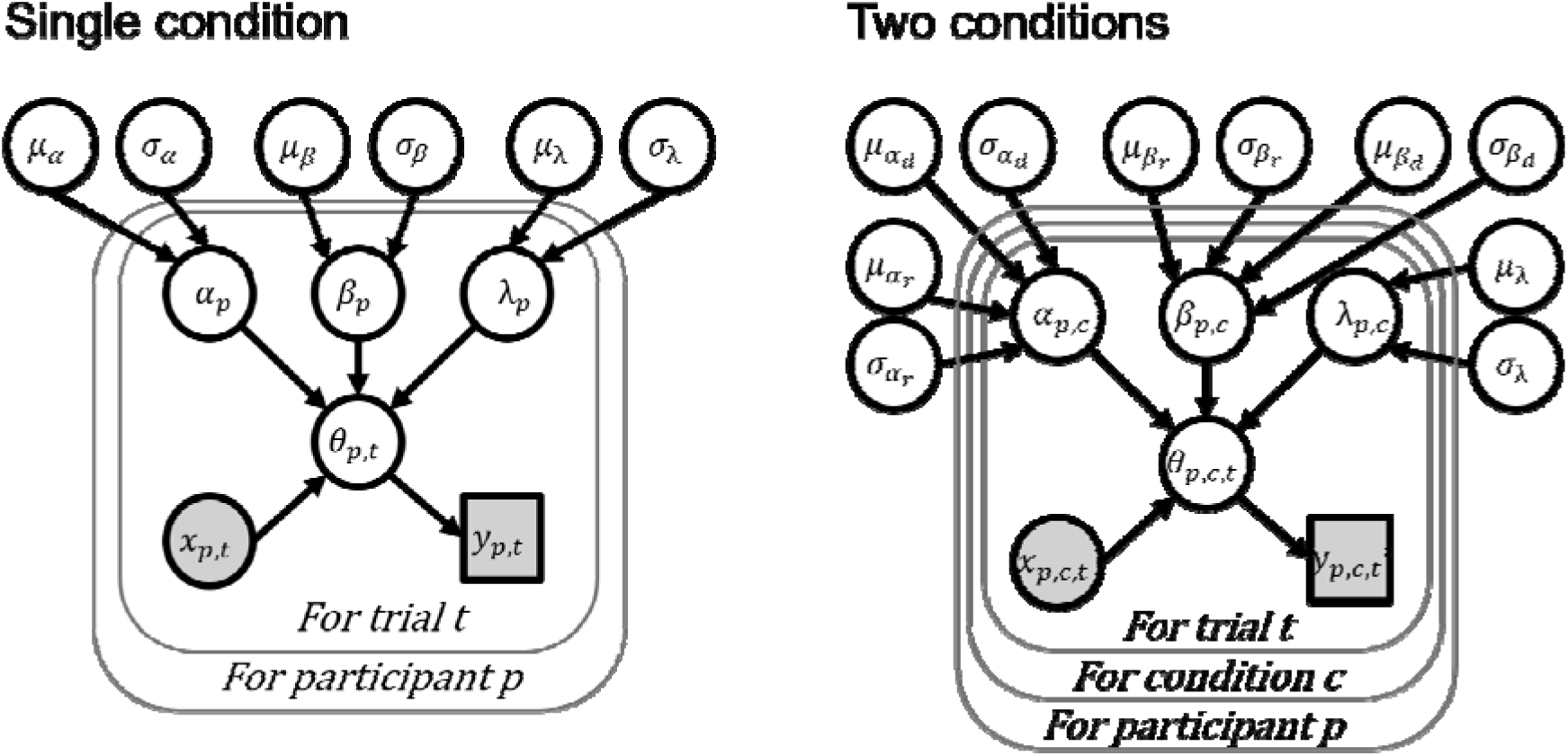
Structure of the hierarchical models for HRDT data. Schematic of the hierarchical models used to estimate psychometric function parameters from trial-wise HRDT data, for either a single condition (left panel) or combined interoceptive and exteroceptive conditions (right panel). Each trial is modelled using stimulus intensity (x) and an observed binary response (y), which is predicted by a psychometric function with parameters specific to each participant and condition. The predicted response probability is denoted by θ. Participant-level parameters, threshold (α), slope (β), and lapse rate (λ), are drawn from group-level distributions with mean (μ) and standard deviation (σ). In the two-condition model, thresholds and slopes are allowed to vary by condition, while the lapse rate is held constant across conditions.

We used weakly informative Gaussian priors for group-level parameters. To constrain slopes to be positive and lapse rates to be in the]0,.5[range, we modelled their distributions across participants in log-space and halved logit-space, respectively.

In the single condition model, the group mean threshold prior (µ_a_) was set so that 95% of its mass spanned from −100 to +100 ΔBPM. Because the impact of the slope parameter f3 depends on the specific PF formulation, we selected priors based on the implied spread of the function, defined as the ΔBPM range between 2.5% and 97.5% response probability, assuming no lapse. Group-level log-slope priors (µ_β_) were chosen such that this spread would fall between 1 and 200 BPM. For the group mean logit-lapse rate parameter (µ_λ_), the prior covered values corresponding to rates ranging from zero to.25 with 95% of its probability mass. Group-level standard deviation parameters (σ_a_,σ_β_,σ_λ_) were given half-Gaussian priors centred at zero, with variances matched to their corresponding group mean priors. A graphical representation of this prior is available in the Supplementary Materials (Fig. S1 and S2)

In the two-condition model, both thresholds and slopes were allowed to vary between the interoceptive and exteroceptive conditions, while the lapse rate was assumed to be equal across conditions. This extension introduced four additional hyperparameters: the group-level difference in threshold (µ_ad_) and log-slope (µ_βd_), and the corresponding between-subject standard deviations (σ_ad_,σ_βd_). Threshold and slope values for the reference condition were indexed with a subscript r. Priors for group differences (µ_ad_,µ_βd_) were Gaussian with mean zero and matched variance to the reference condition priors.

#### RRST model parameterisation

In the RRST, the responses of the participants are transformed to reflect their accuracy (correct/incorrect) rather than the inhalation in which they indicated feeling a higher inspiratory load (first/second). We kept the same accuracy coding for the analysis. These binary responses are modelled using a PF that describes the probability of a correct response as a function of occlusion level (ranging from 0, corresponding to no occlusion, to 17, corresponding to full occlusion). To model this data, we used the Weibull CDF as it is the only PF commonly used for non-negative stimulus intensities (Fig. 1)(Kingdom & Prins, 2010).

The threshold parameter α corresponds to the obstruction level at which participants respond correctly on ∼80% of trials if the lapse rate is zero, marking the Just Noticeable Difference. In conjunction with the threshold α, parameter β determines how rapidly accuracy increases as inspiratory resistance increases. For a constant value of α, higher β values indicate faster transitions from chance to ceiling performance. Of note, this means that parameter β cannot directly serve as an index of the precision of the discrimination process (Kingdom & Prins, 2010). Instead, experimenters seeking to evaluate differences in precision should compare the spread of the psychometric function computed using both α and β. The minimum accuracy is fixed at 50% (chance level) and the upper asymptote of the PF is defined by the lapse rate λ, which accounts for errors due to inattention or reporting mistakes.

We implemented a hierarchical model using the same structure as for the single-condition HRDT (Fig. 2, left panel) but using a Weibull CDF psychometric function. As with the HRDT, the model was implemented in Stan and fitted to trial-wise RRST data.

We used weakly informative Gaussian priors for all group-level parameters. To constrain threshold and slopes to be positive, we modelled their distributions across participants in log-space. Similarly, we modelled the distribution of lapse rates in halved logit-space to ensure that individual participant values were constrained to the]0,.5[range.

For the group mean log-threshold prior (µ_a_), 95% of the prior mass was set to span the full stimulus range (from 1% to 99% occlusion). For the group mean log-slope (µ_β_), we selected a prior corresponding to PF spreads ranging from ∼5% to ∼95% obstruction. The group mean logit-lapse rate prior (µ_λ._) was set to cover rates between zero and.25 with 95% probability. Group-level standard deviations (σ_a_,σ _β_,σ_λ_) were assigned half-Gaussian priors centred at zero, with variances matched to the corresponding group mean priors.

### Parameter and model recovery

We conducted simulation-based validation analyses to assess two aspects of model performance for the HRDT: whether our modelling framework can reliably identify the correct PF formulation (model recovery) and whether it accurately estimates group-level parameters (parameter recovery).

To this end, we generated 100 synthetic datasets for each HRDT PF formulation (Gaussian and Gumbel). Each dataset included 50 simulated participants completing 80 trials of the HRDT (single-condition model). Each synthetic dataset was then fit using both hierarchical Stan models (one per PF formulation). For each model fit, we used four MCMC chains of 2,000 iterations each (1,000 warm-up, 1,000 sampling), with a target acceptance probability of.99 and a maximum tree depth of 12. To reduce computational cost, models with an initialization error, exhibiting divergent transitions after warm-up, or having not properly converged (Rhat>1.05 for any of the parameters) were excluded from further analysis (8 out of 100 for the Gaussian model fitted to Gaussian-generated data, 15 out of 100 for the Gumbel model fitted to Gaussian-generated data, 9 out of 100 for the Gaussian model fitted to Gumbel-generated data; 19 out of 100 for the Gumbel model fitted to Gumbel-generated data; see Table S1 for further details).

Model recovery was evaluated using approximate leave-one-out cross-validation (LOO-CV) with moment matching (Vehtari et al., 2017). For each dataset, we identified the best-fitting model as the one with the highest expected log pointwise predictive density (ELPD). The proportion of correct recoveries (i.e., cases where the model used to generate the data was also selected as the best-fitting model) was tested against chance level (.5) by computing the recovery rate 90% credible intervals (CI) using a Beta(1,1) conjugate prior.

Parameter recovery was evaluated for each group-level means and individual-level values of the PF parameters (threshold, slope, and lapse rate) by computing the proportion of datasets for which the true generating value fell within the 90% posterior credible interval. This was done separately for each model, using only the datasets simulated with the corresponding PF formulation. As with model recovery, we computed 90% CIs for these recovery rates using a Beta(1,1) conjugate prior.

### Model fitting to experimental data

To identify the optimal PF formulation for the HRDT and to derive empirically informed priors for future hierarchical modelling of HRDT and RRST data, we applied our models to the Visceral Mind Project (VMP) dataset.

#### Dataset

The VMP is a large-scale neuroimaging study that was conducted at the Centre of Functionally Integrative Neuroscience (Aarhus University), also described elsewhere (Banellis et al., 2025; Böhme et al., 2025; Ehmsen et al., 2025; Nikolova et al., 2025). A total of 566 participants (360 females, 205 males, one other gender; median age = 24, age range = 18-56) took part in the study. All participants had normal or corrected-to-normal vision, were fluent in Danish or English, and met standard MRI eligibility criteria. Of these, 544 completed at least 60 trials of the HRDT. After excluding 31 participants due to technical errors, data from 513 participants were included in the analysis (324 females, 188 males, one other gender, median age = 24, age range = 18-56). Out of the total sample, 315 participants completed 120 trials of the RRST. Data from 48 were excluded due to suspected equipment malfunction, identified via unusually low task accuracy. The final RRST sample included 267 participants (184 females, 83 males, median age = 24, age range = 18-52).

#### Model formulation and estimation

We applied the hierarchical models described above to the VMP dataset. For the HRDT, each model (one two-condition model for each formulation) was fitted using four MCMC chains of 2,000 iterations per chain, with 1,000 warm-up and 1,000 sampling iterations. The RRST single-condition model struggled to converge and we therefore sampled it using four chains of 6,000 iterations, with 3,000 warm-up and 3,000 sampling iterations. For both HRDT and RRST models, the average target acceptance probability was set to.90 and the maximum tree depth to 12.

All model fits were evaluated using standard convergence diagnostics. We confirmed that no divergent transitions occurred during the sampling phase, that the sampler did not reach the maximum tree depth during that phase, and that posterior exploration was adequate as indicated by energy Bayesian fraction of missing information (E-BFMI) values greater than.1 for all chains. Additionally, all estimated R-hat values were below 1.01 for group-level parameters, consistent with well-mixed chains and reliable posterior estimates, and contraction was larger than.9 for all hyperparameters, indicating that the priors only weakly informed the posterior.

#### Model comparison

To determine which PF provided the best fit for the HRDT data, we performed approximate LOO-CV using the loo package (Vehtari et al., 2017). Model comparison was based on differences in expected log pointwise predictive density (ΔELPD). Differences smaller than four were interpreted as too minor to be of practical significance. For larger ΔELPD values, we considered a model to outperform another when the difference exceeded 1.64 times the standard error of the ΔELPD estimate, corresponding to a conventional one-sided test with a.05 significance threshold. To quantify the strength of evidence, we also calculated the probability that the lower-ranked model would outperform the best-fitting model, assuming Gaussian-distributed ΔELPD estimation errors.

#### Model interpretation

To evaluate condition effects in the HRDT, we examined the posterior distributions of the group-level differences in threshold (µ_ad_) and slope (µ_βd_) between interoceptive and exteroceptive conditions. In practice, we consider that there is sufficient evidence for the existence of a difference when its 95% credible interval does not include zero. Equivalently, we can also compute the posterior probability that the difference is larger or smaller than zero and either use it directly as the test metric, if we have a directional hypothesis about the effect, or to derive a two-sided pseudo p-value using the formula 2 * min(P(θ < 0),1 - P(θ < 0)).

We base inference on posterior probabilities instead of alternative decision metrics, such as Bayes factors, for two reasons. First, probability-based tests map directly onto the interval-based reasoning researchers trained in the frequentist tradition already use, while remaining a genuine statement of belief about the parameter. Second, Bayes factors are sensitive to the scale of the prior under the alternative and thus poorly suited to the weakly informative priors we use here (BARTLETT, 1957).

### Power analysis

When designing experiments, researchers aim to collect a sufficient amount of data to be able to detect an effect if one exists, i.e. have sufficient statistical power. Even though power is a frequentist property, assessing the frequentist properties of Bayesian statistical procedures can be useful to verify their calibration, to guide experimental design, and to have a common metric to compare different Bayesian and non-Bayesian procedures (Gelman et al., 2020; Talts et al., 2018).

To assess the impact of standardised effect size (Cohen’s *d*),key experimental design parameters (namely the number of participants and the number of trials), and analytical choices on this statistical power, we conducted a simulation-based power analysis. The simulated experimental design was analogous to participants completing the interoceptive version of the HRDT under two conditions.

#### Simulations

For each combination of effect size, parameter of interest (slope or threshold), sample size, and trial count, we generated 100 synthetic datasets. We examined effect sizes of 0 (null), 0.2 (small), 0.5 (medium), and 0.8 (large); sample sizes of 15, 30, 60, and 120 participants; and trial counts of 30, 60, and 90 trials per condition.

To build realistic simulated HRD datasets, we simulated synthetic participant responses using the PF shape identified as optimal in the VMP data (i.e., the Gaussian CDF) and we used the □ method to adaptively select stimulus intensities based on these responses, mimicking the functioning of the HRDT toolbox (Legrand et al., 2022). Specifically, in line with the toolbox, we initialized the method with a vector of possible stimulus intensities ranging from-50.5 to +50.5 ΔBPM in 1 BPM increment, the same vector for possible threshold values, and a vector of possible log_10_-slope values ranging from log_10_(0.05) to log_10_(10) in 0.1 increment; we used a uniform prior; we fixed the lapse rate at.02; and the method assumed a cumulative Gaussian PF.

For the control condition, group-level hyperparameters were sampled from the posterior distribution estimated from the VMP dataset. Between-participant standard deviations of the treatment effect were set to half the standard deviation of the corresponding control condition parameter, assuming that between participant variability in the effect of an intervention would be less than baseline between participant variability of the perceptual process. The group mean differences between conditions were set to achieve the specified standardised effect size.

#### Model parameterisation and estimation

Each simulated dataset was fitted using the same hierarchical Bayesian model applied to the VMP dataset. Models were estimated using four MCMC chains with 2,000 iterations each (1,000 warm-up, 1,000 sampling), with a target acceptance probability of.90 and a maximum tree depth of 12. Model fits that failed to meet standard sampling quality criteria, defined as the absence of divergent transitions after warm-up, no iteration exceeding the maximum tree depth, and no chain exhibiting an E-BFMI below.1, were excluded from further analysis.

### Comparison of statistical power under different experimental and analysis designs

Because the true effect size was known for each simulated dataset, we were able to directly assess statistical power, defined as the proportion of simulations in which the null hypothesis (H□) was correctly rejected.

We compared two analytical approaches. In the conventional approach, we used a paired sample t-test to compare the estimates of the psychometric parameters (slope or threshold) returned by the adaptive thresholding algorithm. A p-value below.05 was considered sufficient to reject H□. This corresponds to a scenario where the experimenter extracts the estimates returned by the task script, for each participant and condition, and uses them as dependent variables in subsequent analysis.

In the hierarchical refitting approach, we used a summary of the posterior of our hierarchical model to compare the psychometric parameters across conditions. H□ was rejected if the 95% posterior credible interval for the group mean condition difference did not include zero. This corresponds to a scenario where the experimenter extracts trial level stimulus intensities and binary responses, for each participant and condition, and uses them to fit a hierarchical psychometric function model.

To extrapolate power estimates beyond the specific combinations of effect size, sample size, and trial number used in the simulations, we fitted logistic regression models inStan to predict the probability of H□ rejection as a function of experimental parameters. The model was written as:

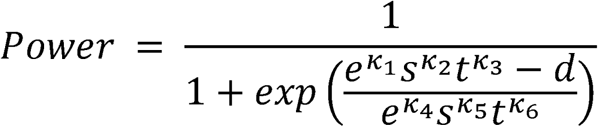

where *d* is the standardised effect size (Cohen’s *d*), *s* is the number of participants, *t* is the number of trials, and the κ terms are regression coefficients estimated from the data. Separate logistic regression models were fit for slope and threshold effects. Each was estimated using the same sampling parameters and diagnostic criteria used for the power analysis fits.

## Results

### Parameter and model recovery

Model comparison successfully selected the generative psychometric function (PF) formulation at rates well above chance (Fig. 3A). Specifically, model recovery rates were 91% (90% CI [.85,.95]) for both models, significantly exceeding the 50% chance level (90% CI did not include.5). These results indicate that our modelling framework is capable of distinguishing between underlying generative functions with high reliability.

**Figure 3.**
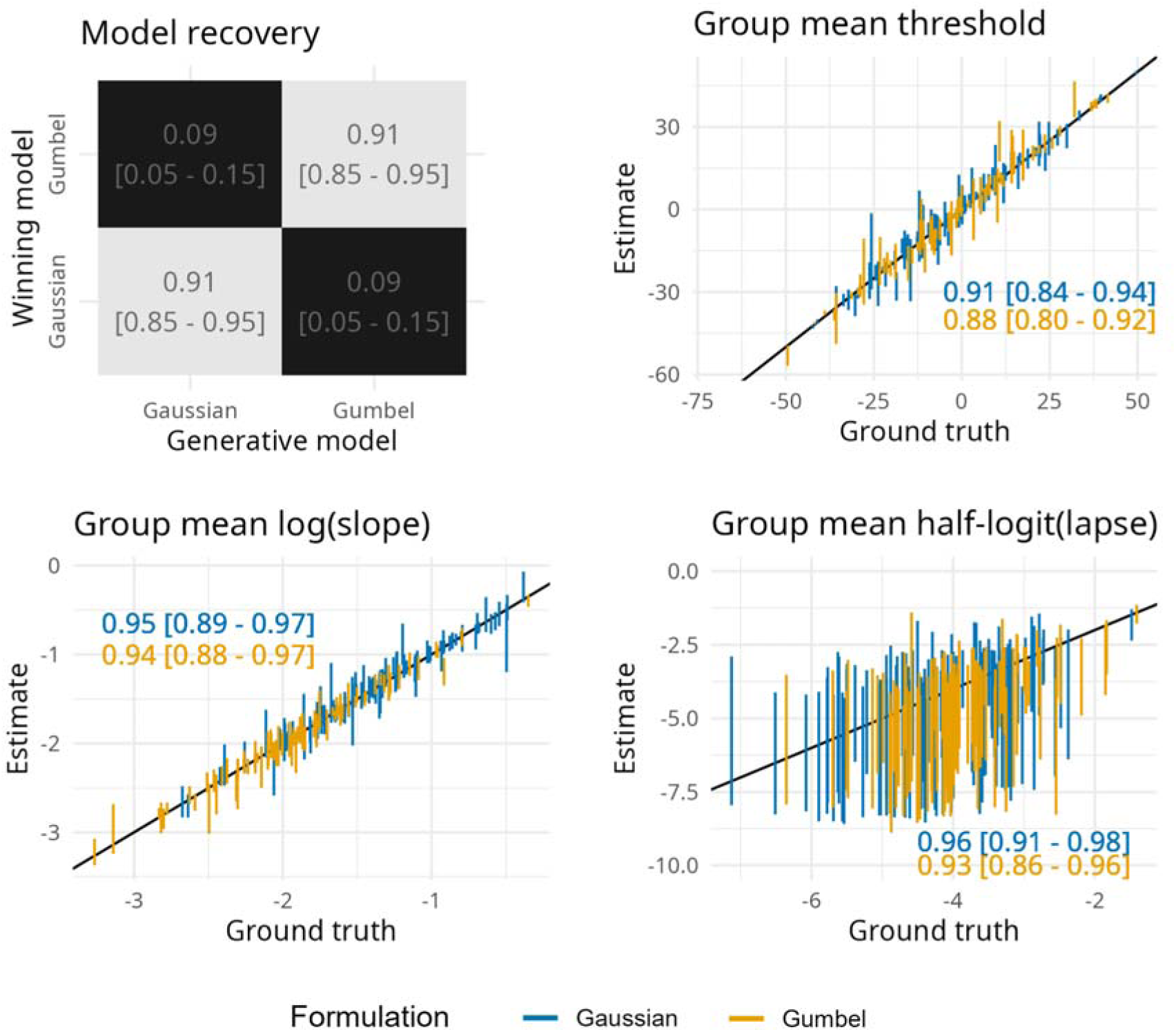
Simulation-based model and parameter recovery analysis. The upper left panel reports model recovery performance, showing the proportion of datasets in which the generating psychometric function (Gaussian or Gumbel) was correctly identified. Recovery rates significantly exceeded chance level (.5), with 90% credible interval (CI) shown in brackets. The remaining panels display parameter recovery results for the group-means: threshold, log(slope), and logit(lapse). Estimated values (y-axis) are plotted against the true generating values (x-axis), with 90% credible intervals represented by vertical bars (blue for Gaussian, yellow for Gumbel). The black identity line indicates perfect correspondence. Recovery rates for each model and parameter are indicated in the lower right of each panel. High recovery rates confirm the reliability of the hierarchical estimation procedure across psychometric function types.

Posterior estimates for group-mean thresholds and slopes closely matched the ground truth values used to simulate the data (Fig. 3B-D). As expected for very low proportions, group mean lapse rates were more difficult to estimate, exhibiting posterior means which often departed from the identity line but with confidence intervals that contained the ground truth value. Since lapse rates are nuisance parameters, exact quantification of their value is not critical. Across all combinations of parameter types and function formulations, parameter recovery rates were high, ranging from 88% (90% CI [.80,.92]) to 96% (90% CI [.91,.98]). In the case of the lapse rate, recovery rates must be interpreted with caution due to high posterior uncertainty.

Similarly, posterior estimates of individual-level threshold and slope parameter values were well aligned with the ground truth values used to simulate the data (Fig. S3), whereas lapse rate estimates were more uncertain. For all parameters, recovery rates were high, ranging from 90% (90% CI [.89,.91]) to 95% (90% CI [.94,.95]).

### Model fitting to experimental data

#### Heart Rate Discrimination Task

Model comparison using the VMP HRDT dataset indicated that the Gaussian PF provided a significantly better fit than the Gumbel formulation (ΔELPD ± SE:-95.8 ± 18.5, P(ΔELPD>0) <.001), consistent with a symmetrical mapping between stimulus intensity (ΔBPM) and response probability.

The fitted Gaussian PF model revealed systematic differences between the interoceptive and exteroceptive conditions. Group-level posterior mean PFs and 95% CI are shown in Fig. 4, together with a representation (shaded areas) of the posterior predictive distribution for individual participants’ PFs, illustrating inter-individual variability.

**Figure 4.**
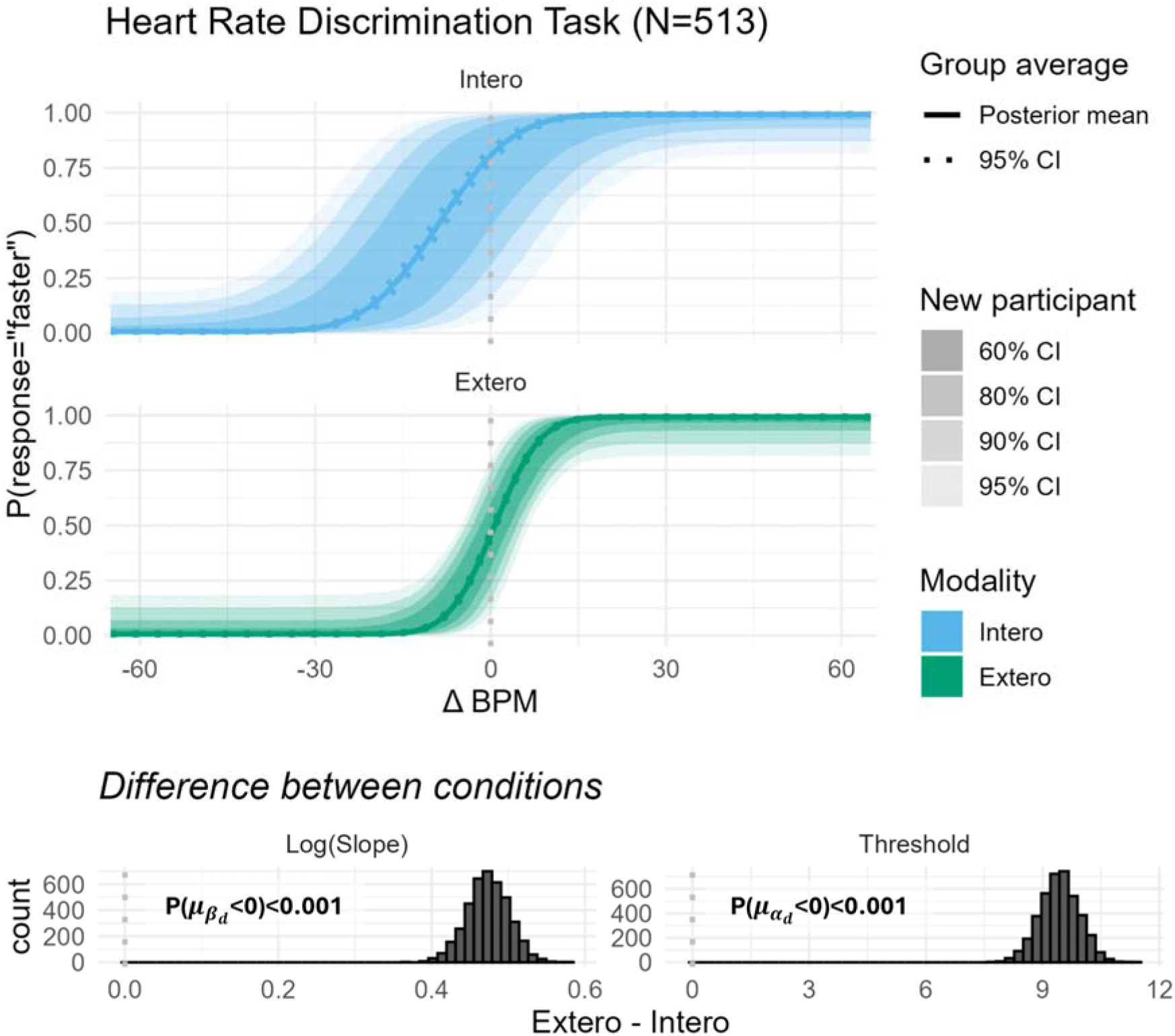
HRDT model fit from the Visceral Mind Project dataset. The top panels display the fitted psychometric functions for the interoceptive (blue) and exteroceptive (green) conditions of the Heart Rate Discrimination Task (N = 513). Solid lines show the group-level posterior mean PFs and dotted lines indicate the 95% credible intervals (CI). Shaded bands represent the expected range of psychometric functions for new participants at varying credible intervals (60%, 80%, 90%, and 95%). The bottom panels show posterior distributions of the group-level differences between conditions for the log-transformed slope (left) and the threshold (right). Distributions are centred well above zero, indicating that interoceptive judgements were both more biased (higher threshold) and less precise (lower slope) compared to exteroceptive judgements. The posterior probabilities for these differences being greater than zero exceed.999, confirming strong evidence for a condition effect.

Posterior distributions of group-level differences between conditions provide clear evidence of these differences. Participants tended to underestimate their heart rate relative to an external auditory cue, as reflected by a positive threshold difference (posterior mean = 9.43, 95% CI [8.44, 10.41]). In addition, interoceptive judgements were less precise than exteroceptive ones, evidenced by a shallower slope (log-slope difference posterior mean = 0.47, 95% CI [0.42, 0.53]). These results indicate that interoceptive performance is characterised by greater bias and reduced reliability compared to exteroceptive comparison judgements.

#### Respiratory Resistance Sensitivity Task

The posterior mean RRST PF and corresponding 95% CI, displayed in Fig. 5, indicate that participants needed high levels of obstruction (∼50%) to start reliably identifying the breath during which resistance was applied. However, most participants reached perfect discrimination at complete obstruction. As for the HRDT plot, inter-individual variability is illustrated by the shaded areas representing the posterior predictive distribution.

**Figure 5.**
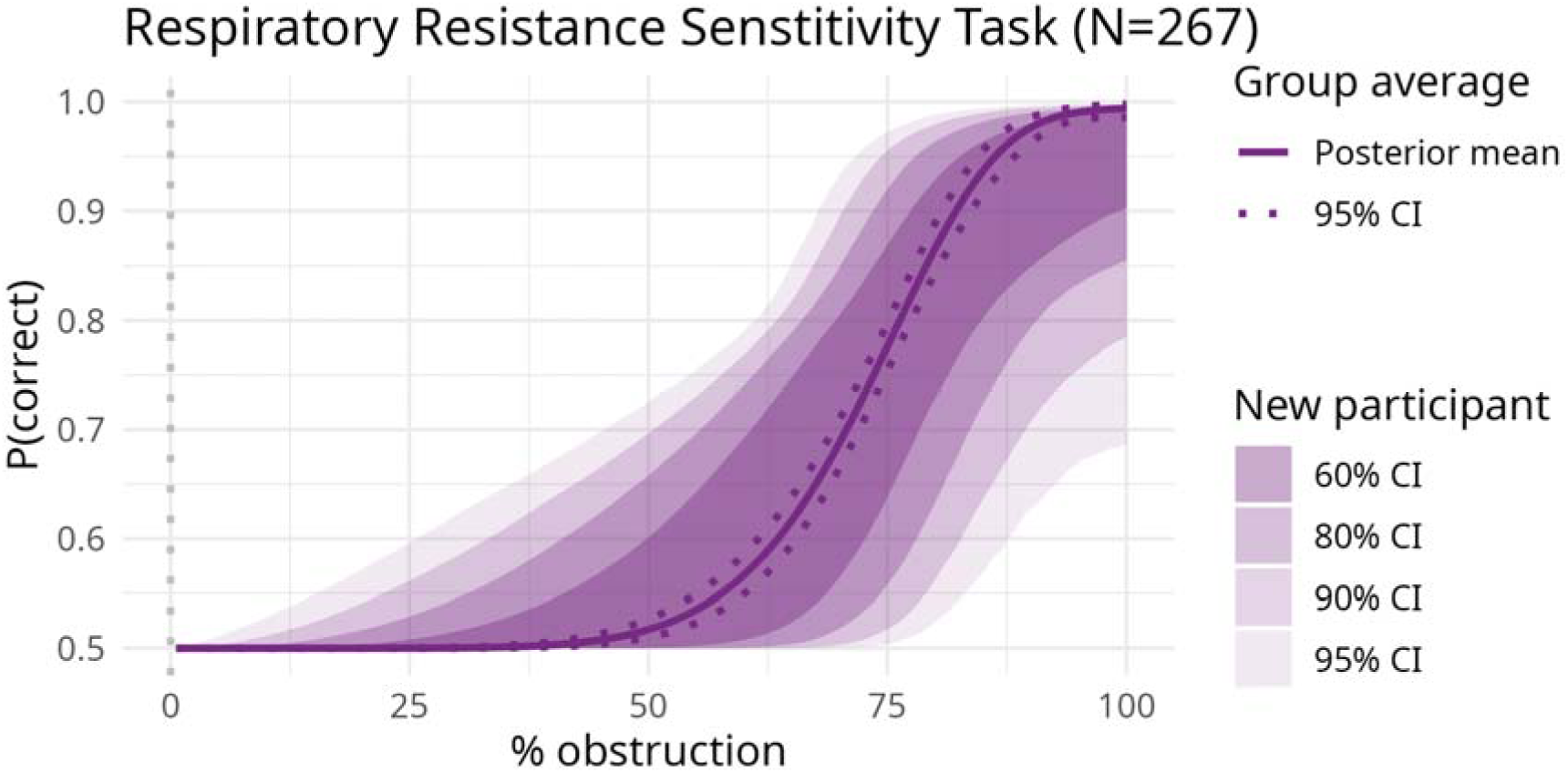
RRST model fit from the Visceral Mind Project dataset. The figure shows the fitted psychometric functions (PF) for the RRST (N = 267). The solid line indicates the group-level posterior mean PF, while the dashed line represents the 95% credible interval. Shaded regions show the posterior predictive range within which a new participant’s psychometric function is expected to fall, visualised across 60%, 80%, 90%, and 95% intervals.

#### Recommended priors

Based on the hierarchical fits to the VMP HRDT and RRST data, we derived empirically informed prior distributions for use in future applications of these models. These recommended priors are implemented in the Stan model files provided in the project repository. To facilitate broader accessibility, we also provide an R script demonstrating how to use the *brms* library to fit HRDT or RRST data using the recommended models and priors; diagnose the fits; and visualise, interpret, and report the results of these fits, without requiring expertise in Stan (Bürkner, 2017).

### Simulation-based power analysis

To evaluate how the number of trials and the number of participants influence our ability to estimate and detect differences in PF parameters (i.e., statistical power), we conducted a simulation-based power analysis. We also aimed to test the theoretical prediction that inference based on hierarchical models would have better power than the common practice of using conventional tests on point estimates returned by the adaptive thresholding algorithm.

Simulation results (Figs. 6 and 7) demonstrate that increasing either the number of participants or the number of task trials consistently increases statistical power. Power was generally higher for threshold than for slope comparisons under matched designs, with threshold comparison power approaching asymptotic levels at lower numbers of trials and participants.

**Figure 6.**
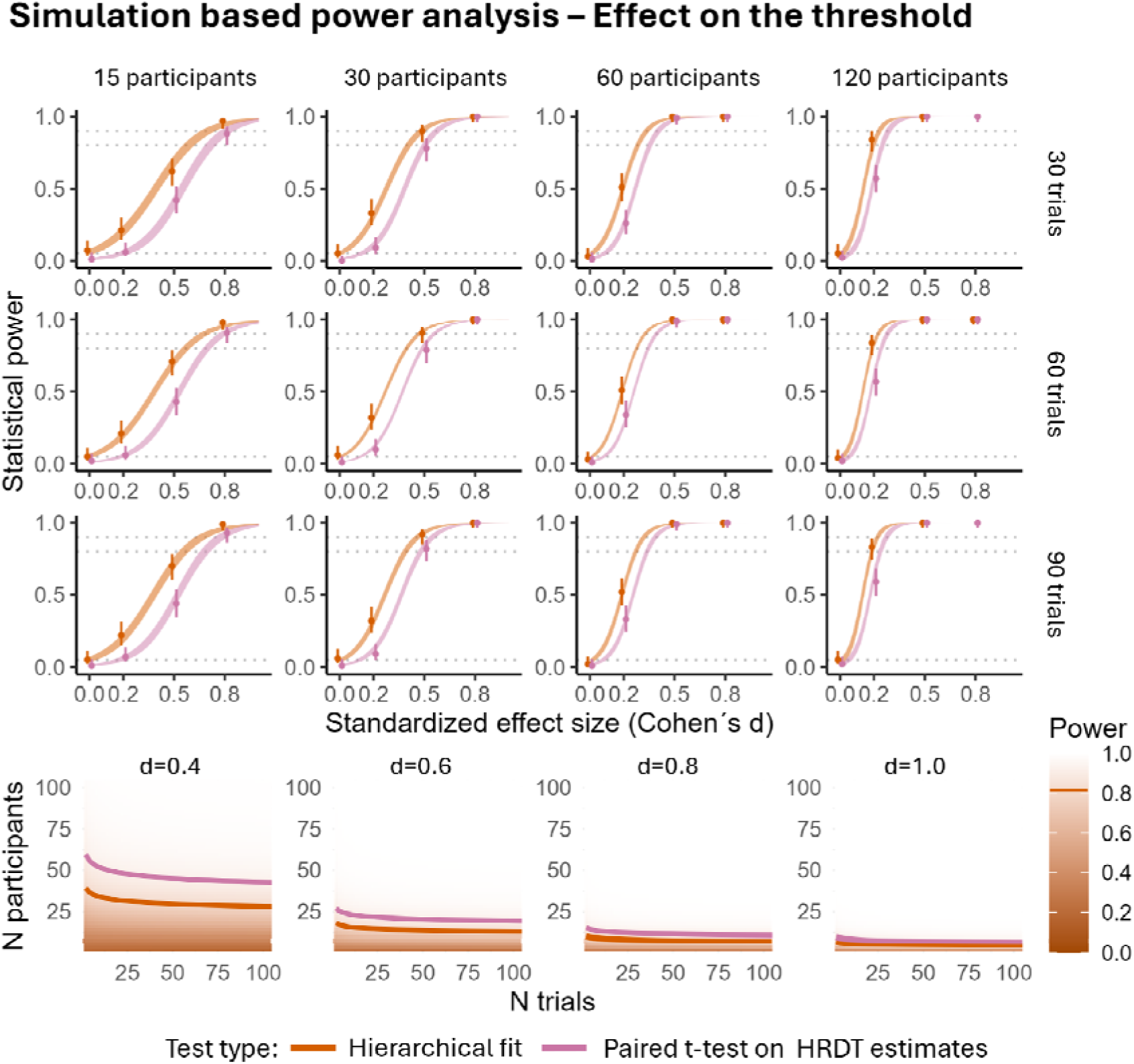
Simulation-based power analysis for a two-session within-participant design: effect on the HRDT threshold. The top panels show simulated power curves for threshold comparisons under varying sample sizes (columns) and trial counts (rows) and across a range of standardised effect sizes (Cohen’s d). Points indicate the observed proportion of simulations in which the null hypothesis was rejected, with vertical lines showing 95% credible intervals. The orange and purple lines correspond to model fits from the hierarchical and paired t-test approaches, respectively. The bottom panels display the power surfaces predicted by the logistic regression model fit to the hierarchical approach simulations. Contour lines indicate combinations of sample size and trial count predicted to yield 80% power for each approach. Hierarchical modelling consistently outperforms the paired sample t-test on extracted L estimates.

**Figure 7.**
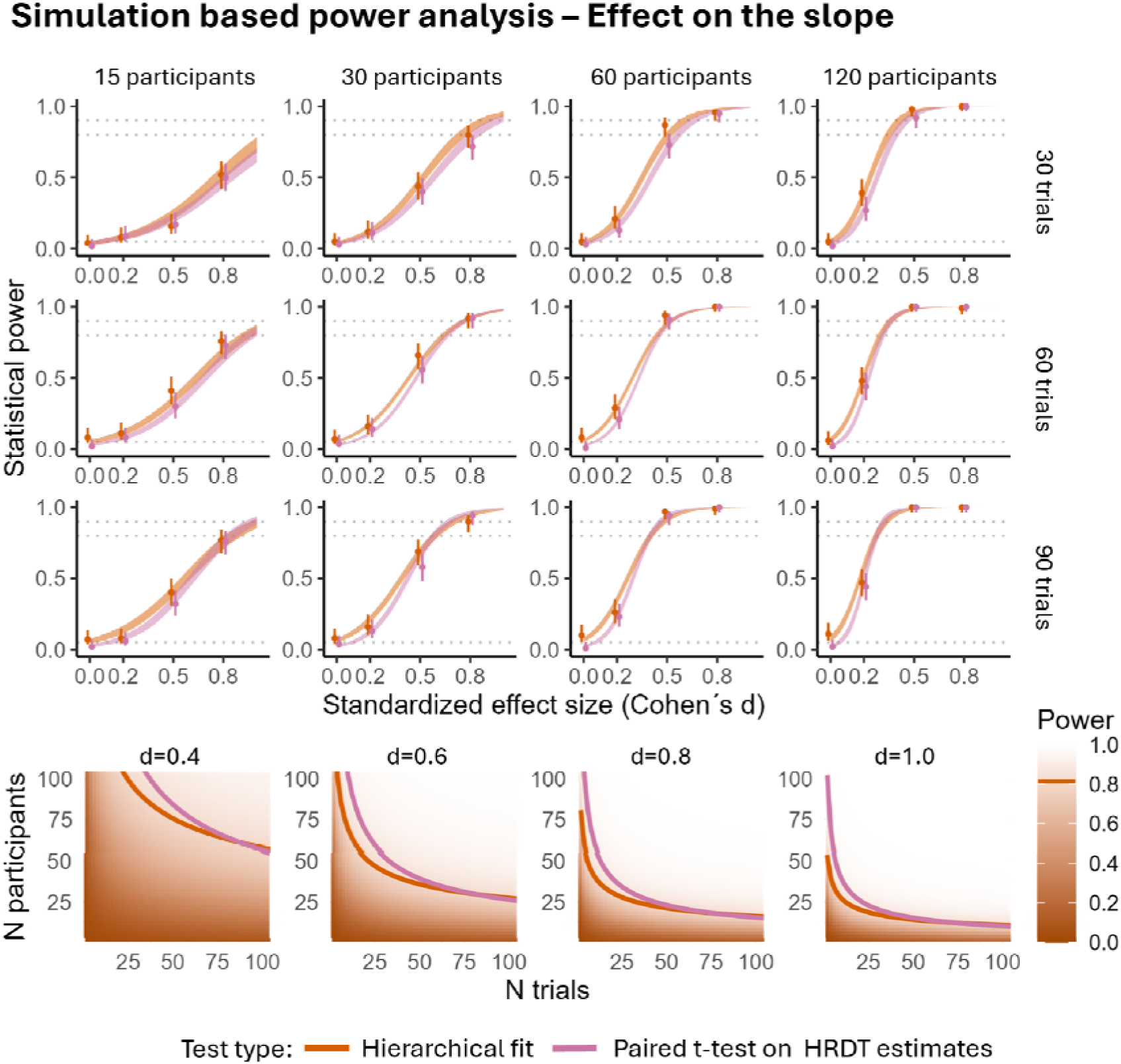
Simulation-based power analysis for a two-session within-participant design: effect on the HRDT slope. The top panels show simulated power curves for slope comparisons under varying sample sizes (columns) and trial counts (rows) and across a range of standardised effect sizes (Cohen’s d). Points indicate the observed proportion of simulations in which the null hypothesis was rejected, with vertical lines showing 95% credible intervals. The orange and purple lines correspond to model fits from the hierarchical and paired t-test approaches, respectively. The bottom panels display the power surfaces predicted by the logistic regression model fit to the hierarchical approach simulations. Contour lines indicate combinations of sample size and trial count predicted to yield 80% power for each approach. Compared to threshold analyses, power increases more gradually, and hierarchical modelling again consistently provides improved sensitivity.

Importantly, comparisons of analytical methods showed that hierarchical Bayesian modelling consistently outperformed the common approach of extracting point estimates from the adaptive thresholding algorithm and analysing them with paired-sample t-tests. This advantage was most pronounced under low-to-moderate sample sizes and effect sizes. For completeness, the Supplementary Materials also include a comparison between our hierarchical modelling approach and a principled use of the paired sample t-test relying on bootstrapping to propagate uncertainty.

Posterior estimates from the logistic regression power models are reported in Table S2 (see Supplementary Materials) and can be used to guide future study design. To facilitate reuse, we provide an interactive Shiny application that allows users to compute statistical power as a function of effect size, number of participants, and number of trials.

## Discussion

This study introduces hierarchical Bayesian modelling of interoceptive performance in the Heart Rate Discrimination Task (HRDT) and Respiratory Resistance Sensitivity Task (RRST). Our initial simulations confirmed that this approach can correctly identify the underlying psychometric function (PF) and accurately estimate both group-and participant-level parameters. We then used it to fit HRDT and RRST data obtained from a large cohort of healthy volunteers. This allowed us to confirm that the Gaussian formulation of the PF provides a good fit to HRDT data and to derive empirically-informed prior distributions for future research. Finally, we used simulations to compare the statistical power achieved by inference based on hierarchical Bayesian models compared to the common practice of running tests on the parameter estimates returned by the HRDT for each participant and condition. These simulations showed that hierarchical Bayesian models consistently achieve superior or equal power to the conventional inference strategy.

Our results demonstrate the practical advantages of hierarchical Bayesian modelling over conventional inference strategies. In addition to being more principled, hierarchical Bayesian models often achieve comparable statistical power with less data. For example, 30 trials per participant across 20 participants are sufficient to detect a standardised threshold difference of 0.5 with 80% power when using hierarchical Bayesian models, whereas the conventional approach would require 30 participants (a 50% increase). Similar benefits are observed for slope estimation: 50 trials per participant across 50 participants is sufficient to detect a 0.5 standardised slope difference with 80% power using hierarchical Bayesian models, while 60 trials are needed with the conventional approach (a 20% increase). These gains in power allow for shorter testing sessions or smaller sample sizes without compromising statistical rigour. While these benefits apply to all researchers working with psychophysical interoception tasks, they are particularly valuable for clinical and paediatric studies, where participant availability and tolerance are often limited.

Importantly, the statistical power gains achieved with hierarchical Bayesian models cannot simply be explained by a difference in the prior information available to the two estimation methods. Indeed, the weakly informative priors used for the hierarchical Bayesian models were less concentrated than the range-uniform priors used to initialize the adaptive thresholding algorithm.

In these analyses, we used weakly informative priors and one may wonder why use priors at all instead of resorting to frequentist alternatives that also support hierarchical structure and partial pooling, such as (generalised) linear mixed-effects models. There are two reasons. First, although the same model structure can in principle be estimated with frequentist methods, in practice such attempts frequently fail to converge, at least with the amounts of data typically available in this field. Second, priors are not merely a technical necessity but a feature we actively embrace: our use of weakly informative priors here reflects only that this was the first dataset collected with these tasks, and one aim of this study was precisely to elicit empirically grounded, informative priors for future researchers working with the HRDT and RRST.

Using such informative priors, both for adaptive thresholding algorithm initialisation and for hierarchical Bayesian model formulation, would likely further improve statistical power: informative priors narrow the plausible range of parameter values, thereby improving stimulus placement and estimation precision. We therefore recommend that future analyses make use of empirically-grounded priors, such as those we derived from the VMP dataset, and we provide updated adaptive thresholding initialisation scripts to this end. Informative priors would additionally enable the principled use of Bayes factors: because Bayes factors are sensitive to the scale of the prior, a justified informative prior renders them interpretable, allowing researchers to quantify evidence for the null as well as for the alternative, a valuable means of distinguishing a genuine absence of effect from insufficient statistical power.

Whereas adopting hierarchical Bayesian modelling has clear benefits, we must also acknowledge some practical limitations. Fitting these models is more computationally intensive than conventional analyses. However, with current software and hardware, this is feasible on standard computers. Moreover, the reduction in data collection time afforded by hierarchical Bayesian models far outweighs the additional analysis time (our demonstration models fit the data of 50 participants in approximately five minutes). Implementing these models also requires some expertise, particularly in selecting appropriate priors and diagnosing model fits. To lower this barrier, we provide an *R Markdown* tutorial (also available as an *html* file) showing how to implement these models, including guidance on model parameterisation, diagnosis, priors, visualisation, and reporting. We chose the *brms* library for this tutorial because it is well-documented, leverages the Stan software for fast and reliable sampling, and uses a syntax similar to popular R regression functions such as *lm*.

We also provide a Shiny app that allows researchers to interactively explore the results of our power analysis. Users can examine how varying numbers of trials, participants, and effect sizes influence statistical power, helping to make informed decisions about study design. Importantly, the app requires no programming experience, making it accessible to a wide range of researchers. While the estimates depend on the specific choices made in our simulations, this represents the first tool for HRDT sample size justification and offers a practical resource for planning future studies.

Beyond interoception, our results are relevant to psychophysics more broadly. This study is among the few to systematically examine how both trial number and participant count jointly influence estimation precision or statistical power in psychophysical designs (Baker et al., 2021; Prins, 2024). We extend these previous studies by systematically assessing statistical power across different effect sizes, highlighting a general asymmetry in the estimation of threshold and slope parameters. Threshold effects are consistently easier to detect than slope effects, with power reaching an asymptote at lower trial and participant numbers. This asymmetry reflects the statistical structure of psychometric functions rather than specific properties of our tasks. In the Gaussian formulation, the threshold corresponds to a mean and the slope (β) to the inverse of a standard deviation (i.e., a precision). Because variance is defined in relation to the mean, its estimation cannot be more precise than that of the mean itself. Although the thresholds and slopes in other PF formulations, like the Weibull used for the RRST, do not map perfectly onto means and variances, they still capture analogous parameters, suggesting that this intuition generalises across PF types. This has direct consequences for experimental design: reliable detection of slope effects will almost invariably require larger samples or trial counts than threshold effects, whereas threshold differences can often be detected with relatively few observations.

Moreover, while our analyses were developed specifically for HRDT and RRST, they can be readily adapted to other tasks, such as those in vision science or thermosensation research. The provided resources offer researchers a straightforward way to implement hierarchical Bayesian approaches in similar paradigms, providing a practical template for extending hierarchical Bayesian modelling across a broad range of psychophysical experiments.

In conclusion, this study presents hierarchical Bayesian models of interoceptive performance in the HRDT and RRST. The approach enables robust estimation of interoceptive bias, sensitivity, and precision directly from trial-wise data. Through simulations and empirical validation, we demonstrate that it improves statistical power over standard analyses, particularly when trial numbers or sample sizes are limited. Our results also provide empirically grounded priors to guide future studies. Accompanied by open-source tools and an interactive power estimation app, this analysis approach supports transparent, efficient, and generalisable model-based inference in interoception research and can readily be adapted to other psychophysical paradigms.

## Declarations

### Ethics approval

The VMP study was performed in line with the principles of the Declaration of Helsinki and was approved by the Central Denmark Region Committees on Health Research Ethics (12.02.20/M-2020-36-20).

### Consent to participate and for publication

All individual participants included in the VMP study gave written informed consent prior to the beginning of testing. This consent included both participation to the experiment and future sharing of the anonymized data.

### Open practices and code and data availability

Neither the VMP experiment (which provided the empirical data) nor the simulation experiments reported in this manuscript were pre-registered. All simulation and analysis scripts, as well as the raw and simulated data used in this manuscript, are available in the project repository: https://github.com/embodied-computation-group/Hierarchical-Interoception.

### Authors’ contributions

Conceptualization: ASC, JFE, MGA; Methodology: ASC, JFE; Formal analysis: ASC, JFE; Resources: MGA; Data Curation: LB, MGA; Writing - Original Draft: ASC; Writing - Review & Editing: ASC, JFE, LB, FF, MGA; Visualization: ASC; Supervision: FF, MGA; Funding acquisition: FF, MGA

