## SupplementaryMaterials for "Hierarchical Bayesian Modelling of Interoceptive Psychophysics"

Supplementary Materials

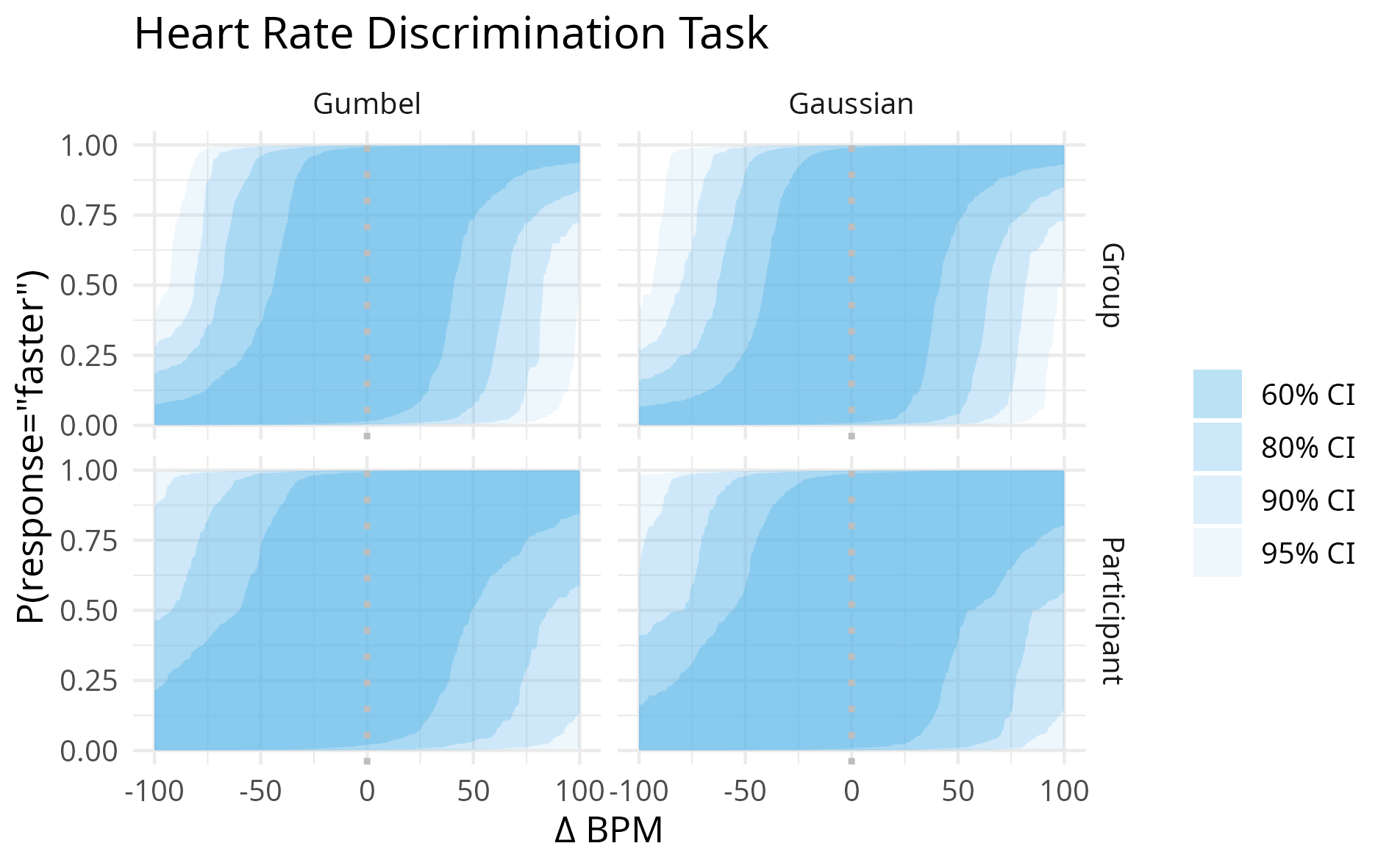

***Figure S1 Prior predictive plots for the HRDT task***

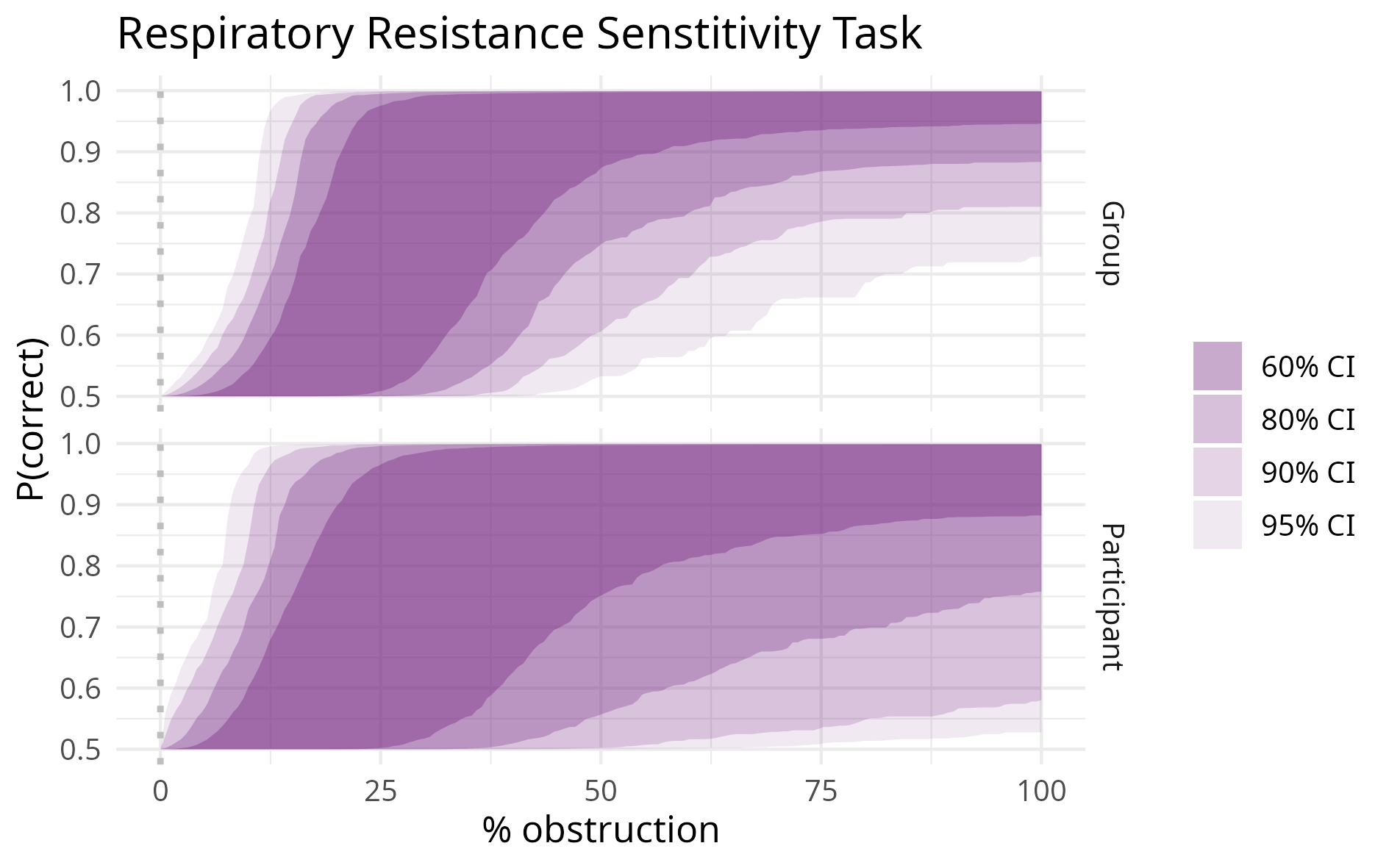

***Figure S2 Prior predictive plots for the RRST task***

| Model type | | Cause(s) of exclusion | | | Number of surviving datasets |
| --- | --- | --- | --- | --- | --- |
| Generative | Fitted | Initialization failure | Divergent transitions | Lack of convergence |  |
| Gaussian | Gaussian | 3 | 4 | 1 | 92 |
| Gaussian | Gumbel | 3 | 12 | 1 | 85 |
| Gumbel | Gaussian | 5 | 4 | 1 | 91 |
| Gumbel | Gumbel | 5 | 14 | 0 | 81 |

***Table S2 Cause for exclusion of model fits from the recovery analyses***

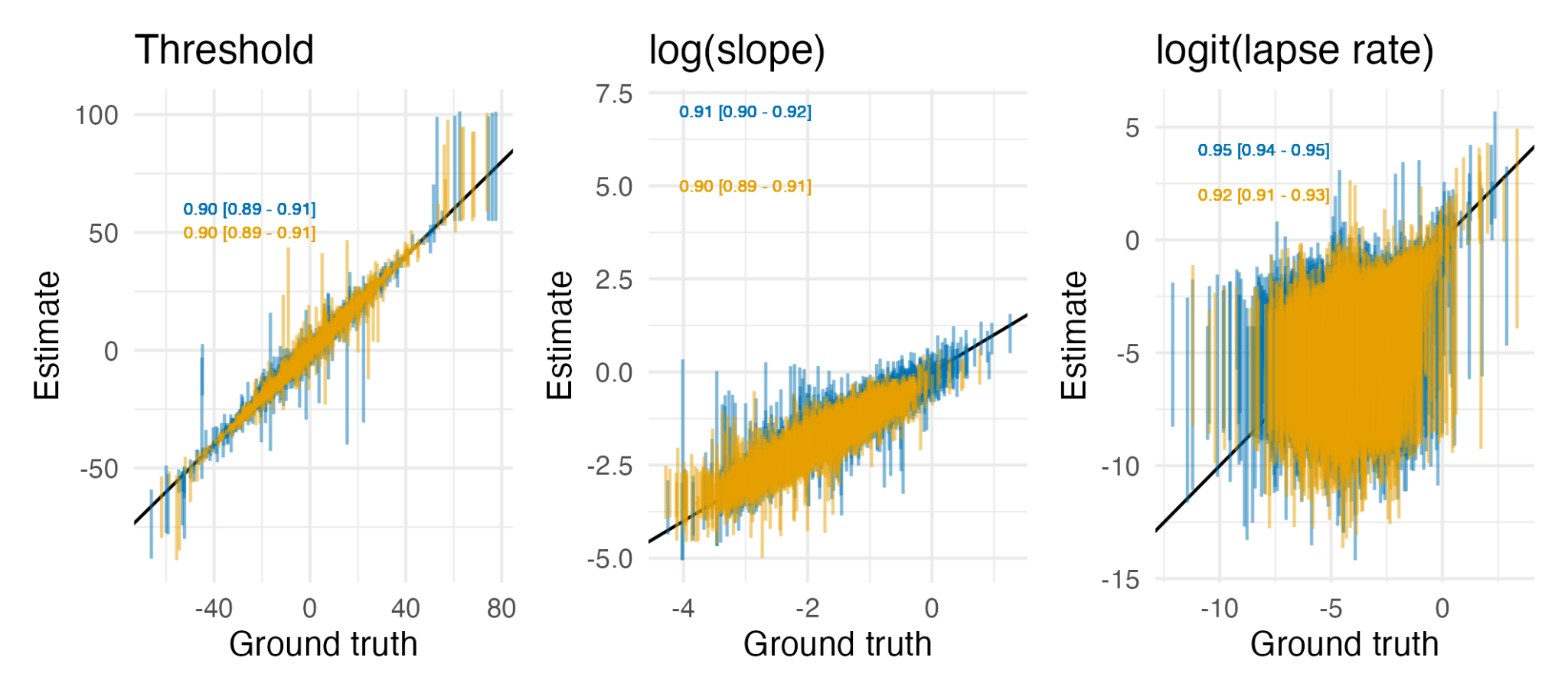

#### Figure S3 Participant level parameter recovery analysis

*Estimated values (y-axis) are plotted against the true generating values (x-axis), with 90% credible intervals represented by vertical bars (blue for Gaussian, yellow for Gumbel). The black identity line indicates perfect correspondence. Recovery rates for each model and parameter are indicated in the lower right of each panel. High recovery rates confirm the reliability of the hierarchical estimation procedure across psychometric function types.*

#### Table S2 *Posterior estimates of the coefficients of the logistic regression model* describing the relationship between statistical power, effect size, number of trials, and sample size for hierarchical models

| **Coefficient** | **Posterior estimates (mean [95%CI])** | | | |
| --- | --- | --- | --- | --- |
|  | **Hierarchical fit** | | **HRDT estimates** | |
|  | **Threshold** | **Slope** | **Threshold** | **Slope** |
| $\kappa_{1}$ | 0.51 [0.18 ; 0.85] | 2.16 [1.86 ; 2.47] | 0.95 [0.71 ; 1.19] | 2.20 [1.93 ; 2.47] |
| $\kappa_{2}$ | -0.50 [-0.54 ; -0.46] | -0.57 [-0.61 ; -0.52] | -0.52 [-0.55 ; -0.49] | -0.53 [-0.56 ; -0.49] |
| $\kappa_{3}$ | -0.03 [-0.10 ; 0.05] | -0.26 [-0.33 ; -0.20] | -0.04 [-0.10 ; 0.01] | -0.28 [-0.34 ; -0.23] |
| $\kappa_{4}$ | -0.19 [-0.76 ; 0.39] | 0.33 [-0.25 ; 0.95] | -0.73 [-1.36 ; -0.10] | 1.51 [0.88 ; 2.15] |
| $\kappa_{5}$ | -0.57 [-0.65 ; -0.49] | -0.52 [-0.60 ; -0.43] | -0.48 [-0.57 ; -0.39] | -0.57 [-0.65 ; -0.48] |
| $\kappa_{6}$ | -0.07 [-0.20 ; 0.05] | -0.11 [-0.24 ; 0.01] | -0.03 [-0.17 ; 0.11] | -0.39 [-0.52 ; -0.26] |

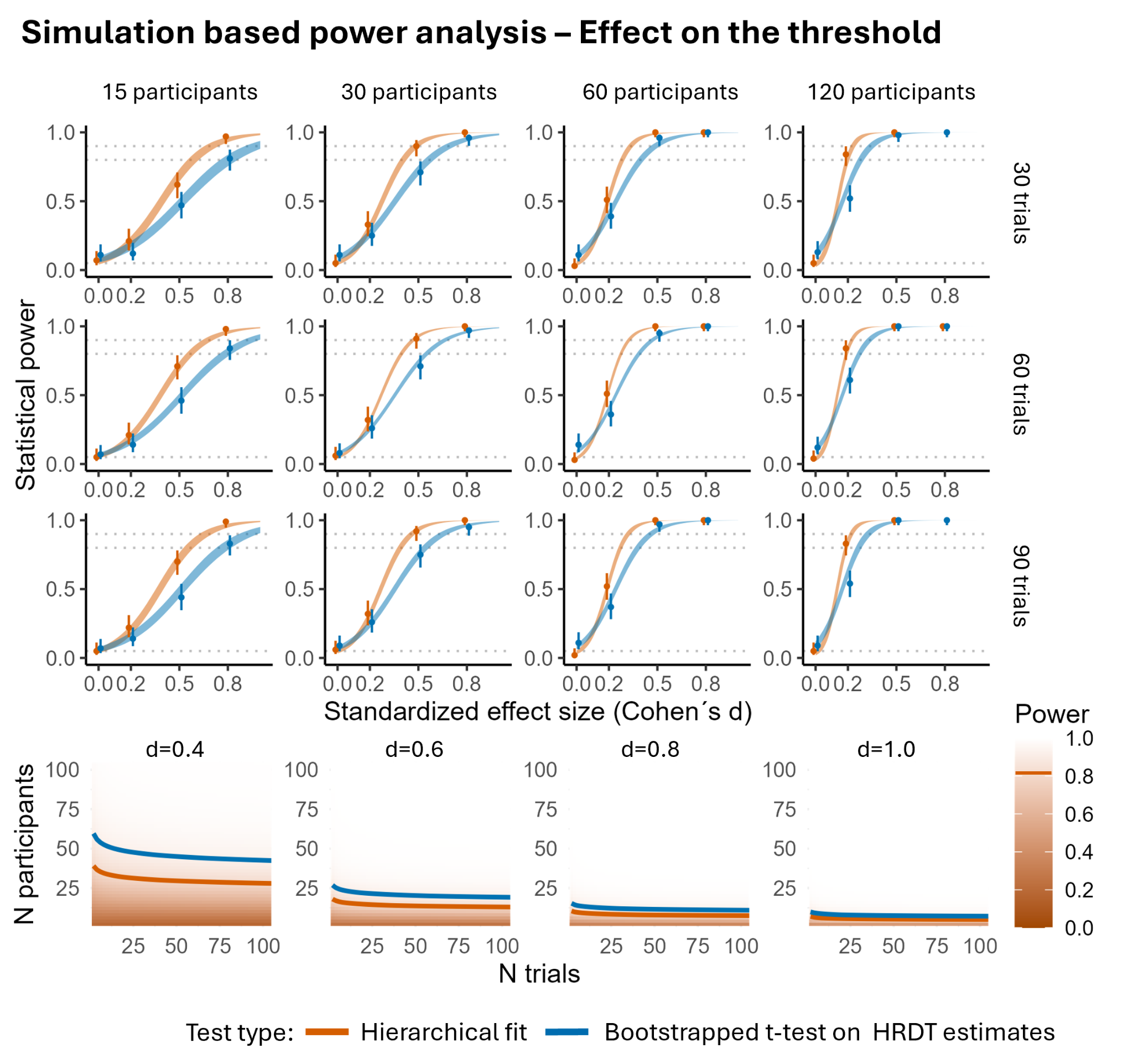

#### Figure S4 Comparison of statistical power achieved by inference based on hierarchical models compared to a bootstrapped paired sample t-test: effects on the HRDT threshold

*The top panels show simulated power curves for threshold comparisons under varying sample sizes (columns) and trial counts (rows) and across a range of standardised effect sizes (Cohen’s d). Points indicate the observed proportion of simulations in which the null hypothesis was rejected, with vertical lines showing 95% credible intervals. The orange and blue lines correspond to model fits from the hierarchical and bootstrapped paired t-test approaches, respectively. The bottom panels display the power surfaces predicted by the logistic regression model fit to the hierarchical approach simulations. Contour lines indicate combinations of sample size and trial count predicted to yield 80% power for each approach. Hierarchical modelling consistently outperforms the paired sample t-test on extracted 𝜓 estimates.*

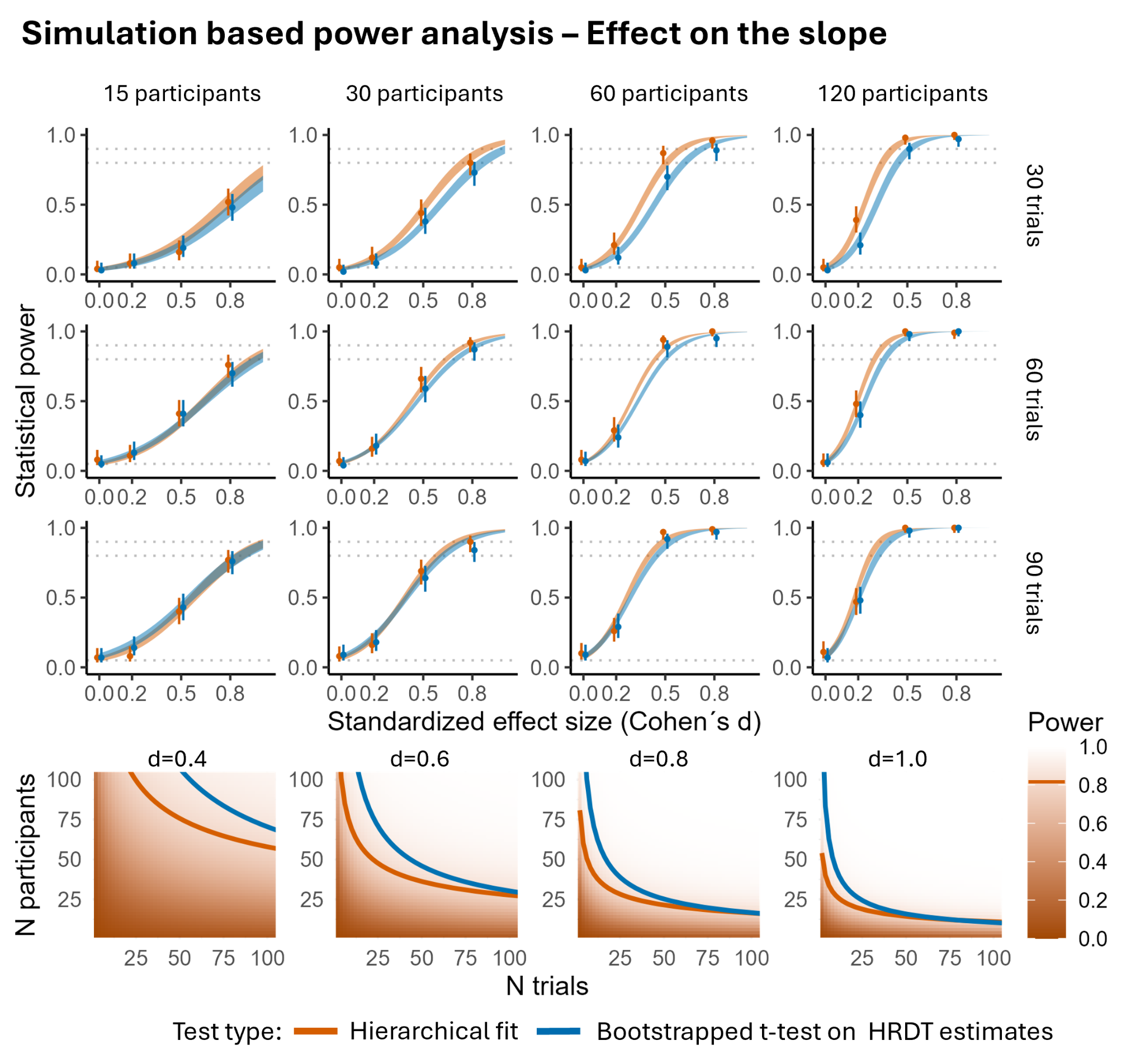

#### Figure S5 Comparison of statistical power achieved by inference based on hierarchical models compared to a bootstrapped paired sample t-test: effects on the HRDT slope

*The top panels show simulated power curves for threshold comparisons under varying sample sizes (columns) and trial counts (rows) and across a range of standardised effect sizes (Cohen’s d). Points indicate the observed proportion of simulations in which the null hypothesis was rejected, with vertical lines showing 95% credible intervals. The orange and blue lines correspond to model fits from the hierarchical and bootstrapped paired t-test approaches, respectively. The bottom panels display the power surfaces predicted by the logistic regression model fit to the hierarchical approach simulations. Contour lines indicate combinations of sample size and trial count predicted to yield 80% power for each approach. Hierarchical modelling consistently outperforms the paired sample t-test on extracted 𝜓 estimates.*

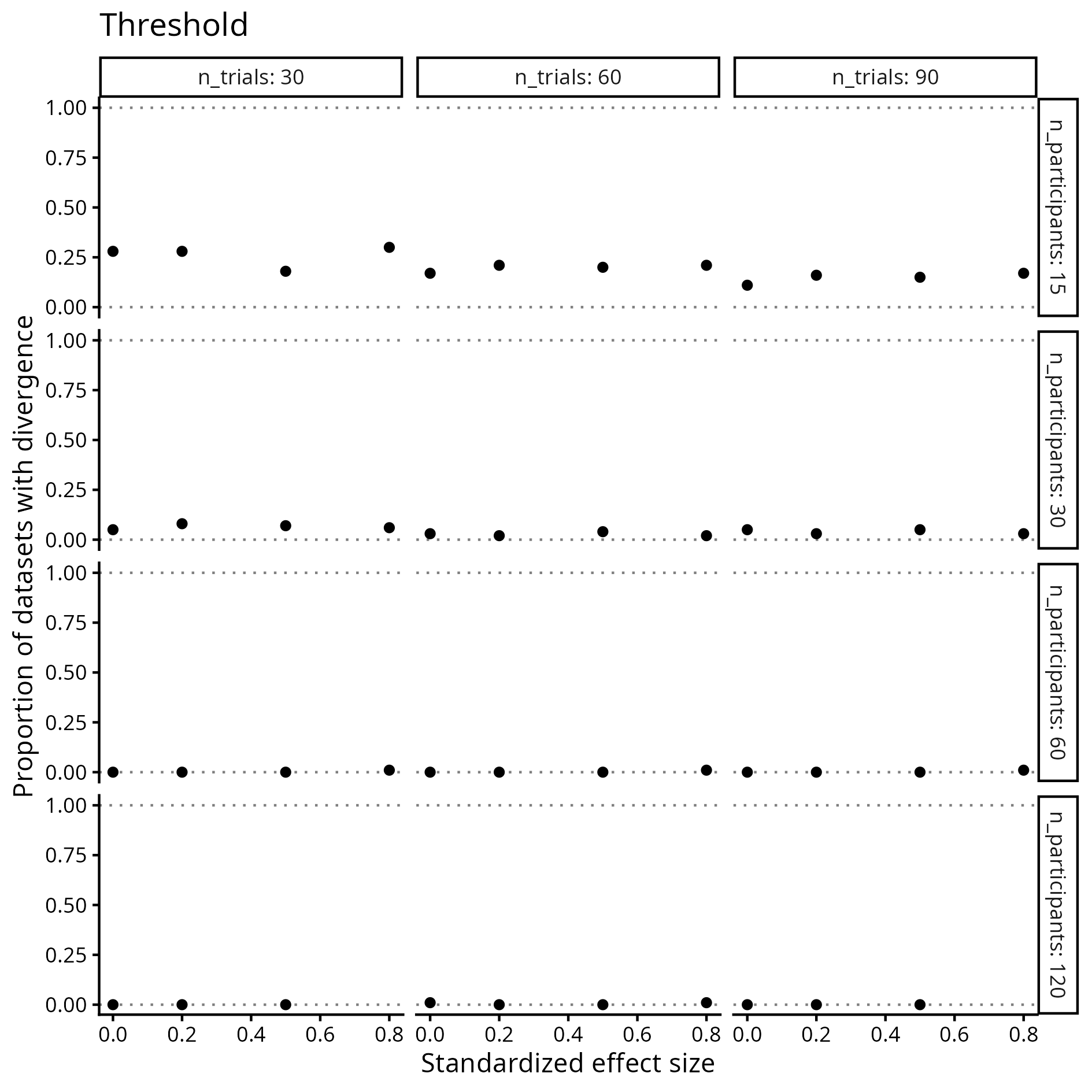

#### Figure S6 Proportion of simulated datasets rejected from the threshold power analysis because the hierarchical model fit contained divergent transitions

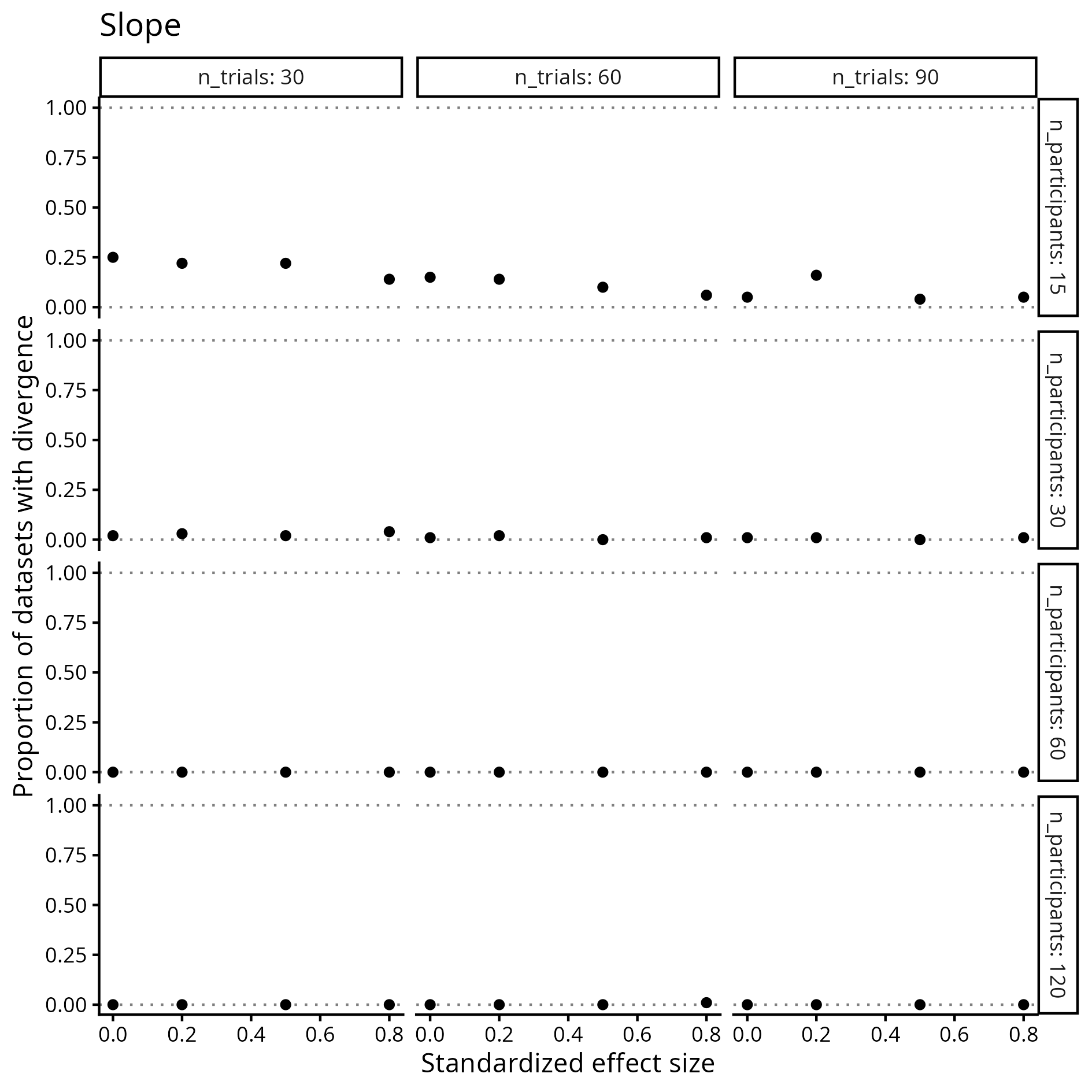

#### Figure S7 Proportion of simulated datasets rejected from the slope power analysis because the hierarchical model fit contained divergent transitions
